# Experimental Determination of Evolutionary Barriers to Horizontal Gene Transfer

**DOI:** 10.1101/722959

**Authors:** Hande Acar Kirit, Mato Lagator, Jonathan P. Bollback

## Abstract

Horizontal gene transfer, the acquisition of genes across species boundaries, is a major source of novel phenotypes. Several barriers have been suggested to impede the likelihood of horizontal transmission; however experimental evidence is scarce. We measured the fitness effects of genes transferred from *Salmonella enterica* serovar Typhimurium to *Escherichia coli*, and found that most result in strong fitness costs. Previously identified evolutionary barriers — gene function and the number of protein-protein interactions — did not predict the fitness effects of transferred genes. In contrast, dosage sensitivity, gene length, and the intrinsic protein disorder significantly impact the likelihood of a successful horizontal transfer. While computational approaches have been successful in describing long-term barriers to horizontal gene transfer, our experimental results identified previously underappreciated barriers that determine the fitness effects of newly transferred genes, and hence their short-term eco-evolutionary dynamics.

## Introduction

Horizontal gene transfer (HGT) is the lateral transfer of genetic material between different individuals and species, and a major driver of evolution in all domains of life, but particularly in bacteria (Ochman, Lawrence, & Groisman, 2000; Polz, Alm, & Hanage, 2013; Popa & Dagan, 2011). HGT has contributed to the stunning array of phenotypic diversity we observe in nature. As a source of phenotypic novelty, HGT differs from *de novo* mutations as new adaptive phenotypes can appear faster and sweep through the populations more rapidly. Among these novelties, some HGT events even have profound impact on our daily lives. For instance, the mannanase gene of the coffee berry borer beetle, *Hypothenemus hampei*, was acquired from bacteria and it enables the beetle to digest the complex sugars in coffee beans, thus turning the beetle into an industrially relevant pest (Acuña et al., 2012). Similarly, the horizontal acquisition of vancomycin resistance by *Staphylococcus aureus* (Weigel et al., 2003) and virulence factors by *Escherichia coli* O104:H4 in the 2011 European outbreak (Rasko et al., 2011) are serious public health concerns. Despite the importance of HGT, we still have a limited understanding of the factors that determine the outcome of HGT events.

The success, or failure, of an HGT event is largely determined by the effect of the transferred gene on the fitness of the recipient species. Deleterious genes will be purged by selection, beneficial genes may be fixed, and the fate of neutral genes will be determined by stochastic forces, such as genetic drift or genetic draft (Gillespie, 2000; Soucy, Huang, & Gogarten, 2015). Thus, the fate of transferred genes depends on the distribution of their fitness effects (DFE). To date, we have little knowledge of the DFE for newly transferred genes, or of the factors that determine those fitness effects, especially following expression in the recipient cell.

Motivated by computational analyses, several non-mutually exclusive factors, or ‘selective barriers’, have been hypothesized to affect the DFE of horizontally transferred genes: the functional category of the transferred genes (Jain, Rivera, & Lake, 1999; Rivera, Jain, Moore, & Lake, 1998), the number of protein-protein interactions (PPI) (Cohen, Gophna, & Pupko, 2011; Jain et al., 1999), and the divergence between the donor and the recipient cell that is inferred as a difference in their GC content or the codon usage bias (Baltrus, 2013; Drummond & Wilke, 2009; Tuller et al., 2011). While computational approaches have produced valuable insights into HGT, there are important barriers only assessable through experimental analyses. Notably, the suggestion that gene dosage, or dosage sensitivity, might play a role in HGT came from experimental insights and cannot be addressed with sequence data alone (Papp, Pál, & Hurst, 2003; Sorek et al., 2007).

Here we conduct a systematic experimental test of the barriers to HGT by transferring and expressing orthologs from *Salmonella enterica* serovar Typhimurium to *Escherichia coli*, and measuring their fitness effects. Specifically, we asked: What is the DFE of newly transferred genes? What selective barriers affect the likelihood of a newly transferred gene’s spread in a population?

## Results

We mimicked HGT by transferring genes from *S*. Typhimurium to an *E. coli* recipient to determine the DFE and test whether selective barriers — functional category, number of PPI, GC content, codon usage, and dosage sensitivity — affect the likelihood of transfer. We chose these species as they share the same environment – mammalian gut – which increases the potential of HGT between them. *S*. Typhimurium and *E. coli* are also close relatives (McClelland et al., 2001). This similarity means that most of the native PPIs of the transferred orthologs are likely to be preserved. We transferred a total of 44 genes, placed them under the control of the same promoter, induced their expression, and measured their fitness effects with competition assays (Fig. 1). We verified the expression of a subset of transferred genes at the level of translation (Sup. Fig. 1). The recipient *E. coli* strain was chromosomally labeled with two different fluorescent markers: CFP-labeled ‘mutant’ strain carried the plasmid containing one of the introduced genes (Sup. Fig. 2); and YFP-labeled ‘wild-type’ strain carried the same plasmid only without the introduced gene. Two strains were mixed at equal frequencies and grown together with samples taken at regular intervals, and the change in frequencies of the two strains were measured by flow cytometry to determine fitness. The fitness effects, or the selection coefficients (*s*), of transferred genes were estimated using the regression model *ln*(1+*s*) = (*ln*R_t_ – *ln*R_0_)/t, where R is the ratio of mutant to wild type, and t is the number of generations (Elena, Ekunwe, Hajela, Oden, & Lenski, 1998). By performing 32 replicates for each transferred gene, we estimated the fitness effects of transferred genes with a previously not achievable degree of precision, Δ*s* ≈ 0.002.

**Fig. 1.**
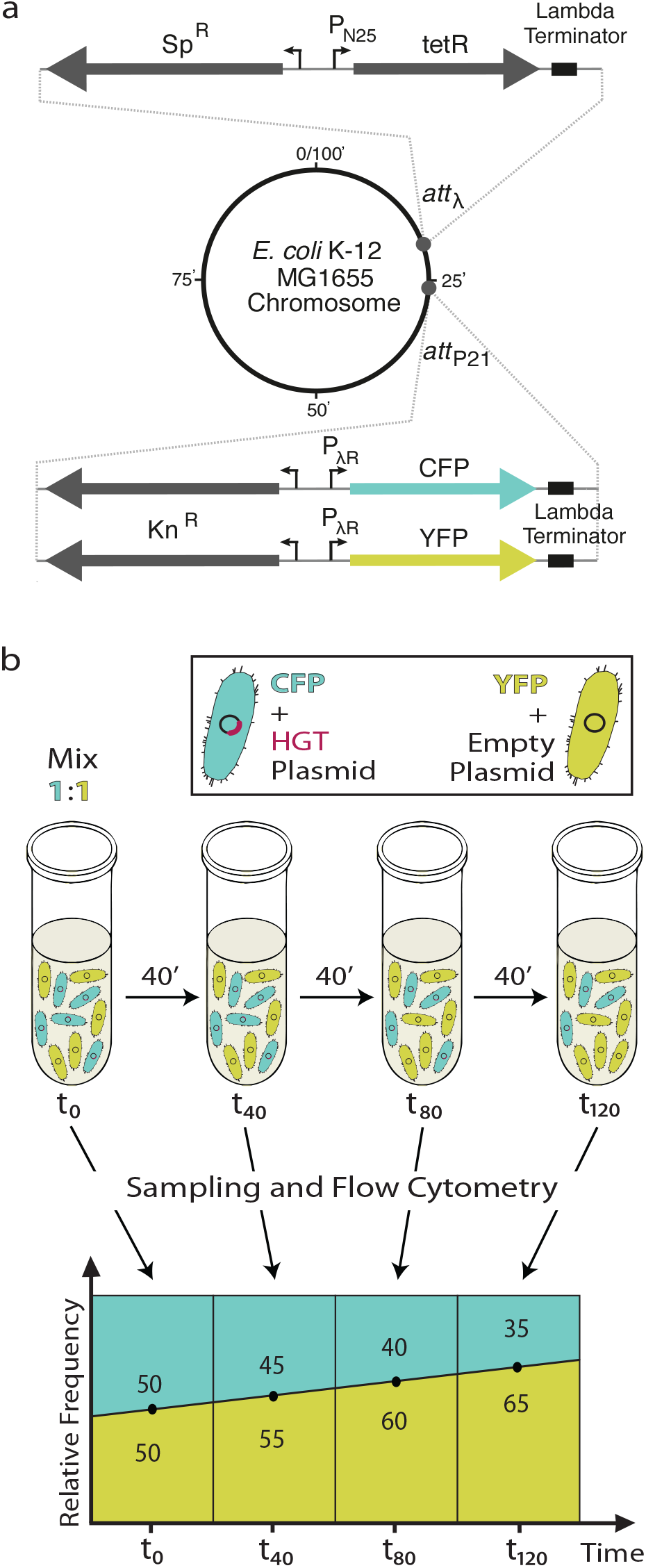
Schematic representation of the experimental design. **a**. Chromosomal modifications in the recipient strain, E. coli MG1655 att-λ::(tetR-Sp^R^) att-p21::(CFP/YFP-Kn^R^), for the transferred genes. att-λ and att-p21correspond to the attachment sites of the phage λ and p21, respectively. TetR is the repressor protein controlling the expression of the transferred genes. Sp^R^ and Kn^R^ are the resistance genes for spectinomycin and kanamycin, respectively. P_N25_ and P_λR_ are the constitutive promoters. See Materials and methods section for details. **b**. Depiction of the competition assay. Blue cells with CFP represent the ‘mutant’ strain that carries the pZS*-HGT plasmid containing the introduced gene, whereas yellow cells with YFP represent the ‘wild type’ strain that carries the empty pZS*-HGT plasmid without the introduced gene. The plot illustrates an example where the fitness effect of the gene is deleterious, resulting in a decrease in the frequency of blue cells over time. Numbers inside the segments represents the frequency of the type of the cell with same color.

### Distribution of fitness effects

Majority of *S*. Typhimurium genes transferred into *E.coli* had a negative effect on fitness with a median fitness effect *s* = – 0.020 and a range of – 0.606 to 0.009 (Fig. 2). Out of 44 transferred genes, 3 were beneficial, 5 were neutral (fitness not significantly different from zero), 25 were moderately deleterious, and 11 were highly deleterious (s < – 0.1, Sup. Table 1). The DFE shares the general features with experimentally determined DFEs for other biological systems, such as mutations in bacterial promoters (Kinney, Murugan, Callan, & Cox, 2010), viral sequences (Sanjuán, Moya, & Elena, 2004), transcription factors (Shultzaberger, Maerkl, Kirsch, & Eisen, 2012), and random transposon insertions (Elena et al., 1998). The effect of the deleterious transferred genes is well described by a log-normal distribution (μ = −3.562 and σ = 1.693), as previously observed for DMEs of mutations (Eyre-Walker & Keightley, 2007).

**Fig. 2.**
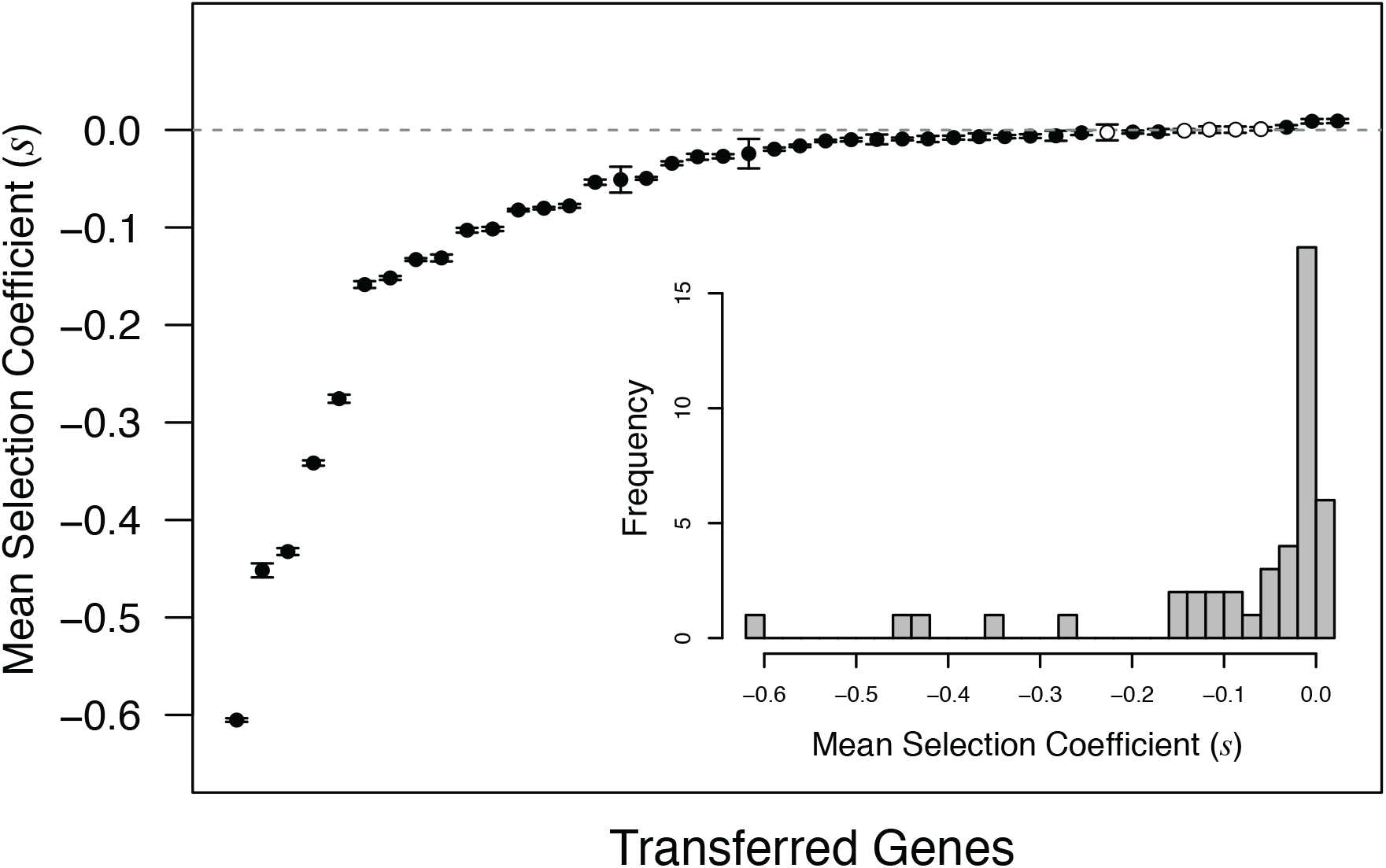
DFE of 44 newly transferred S. Typhimurium orthologs expressed in E. coli. On the x-axis, genes are sorted by their fitness effects (s). Error bars indicate the 95% CI of the selection coefficients from 32 replicate measurements for each gene. Empty circles represent the 5 genes with effects not significantly different from zero. Embedded plot gives the histogram representation of the same data.

### Evaluating potential barriers to HGT

The 44 genes chosen for experimental transfer differ with respect to several factors hypothesized to act as barriers to HGT. We asked whether these selective barriers predicted the fitness effects of the transferred genes. In particular, we looked at the effect of functional category, number of PPIs, gene length, and deviation in the GC content and codon usage between the transferred S. Typhimurium genes and their orthologs in E. coli (Sup. Table 2). We tested for these factors in a multiple linear regression model framework (F_5,37_ = 2.24, Sup. Table 3, for details see Materials and Methods – Statistical analyses). Surprisingly, we found a significant effect only for the gene length.

### Functional gene category

The ‘functional category hypothesis’ proposes that informational genes (those involved in DNA replication and repair, transcription, and translation) are less amenable to transfer than operational genes (those involved in processes like metabolism and biosynthesis) (Rivera et al., 1998; Jain et al., 1999; Nakamura, Itoh, Matsuda, & Gojobori, 2004). To test this hypothesis, we grouped genes by Gene Ontology and MultiFun annotations (Karp et al., 2007), classifying 24 genes as informational genes and 19 as operational. No significant difference was observed between the two categories in either mean fitness effects (Fig. 3a, Wilcoxon rank sum test: median ***s_Info_*** = −0.026, median ***s_Oper_***= −0.010, *W* = 181, *p* = 0.130; multiple regression model: *p* = 0.354) or in the variance in fitness effects (Levene’s test: 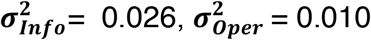, *F_1,41_* = 1.164, *p* = 0.287). Interestingly, 4 of the 5 most deleterious genes 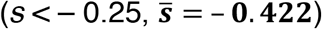 were informational.

**Fig. 3.**
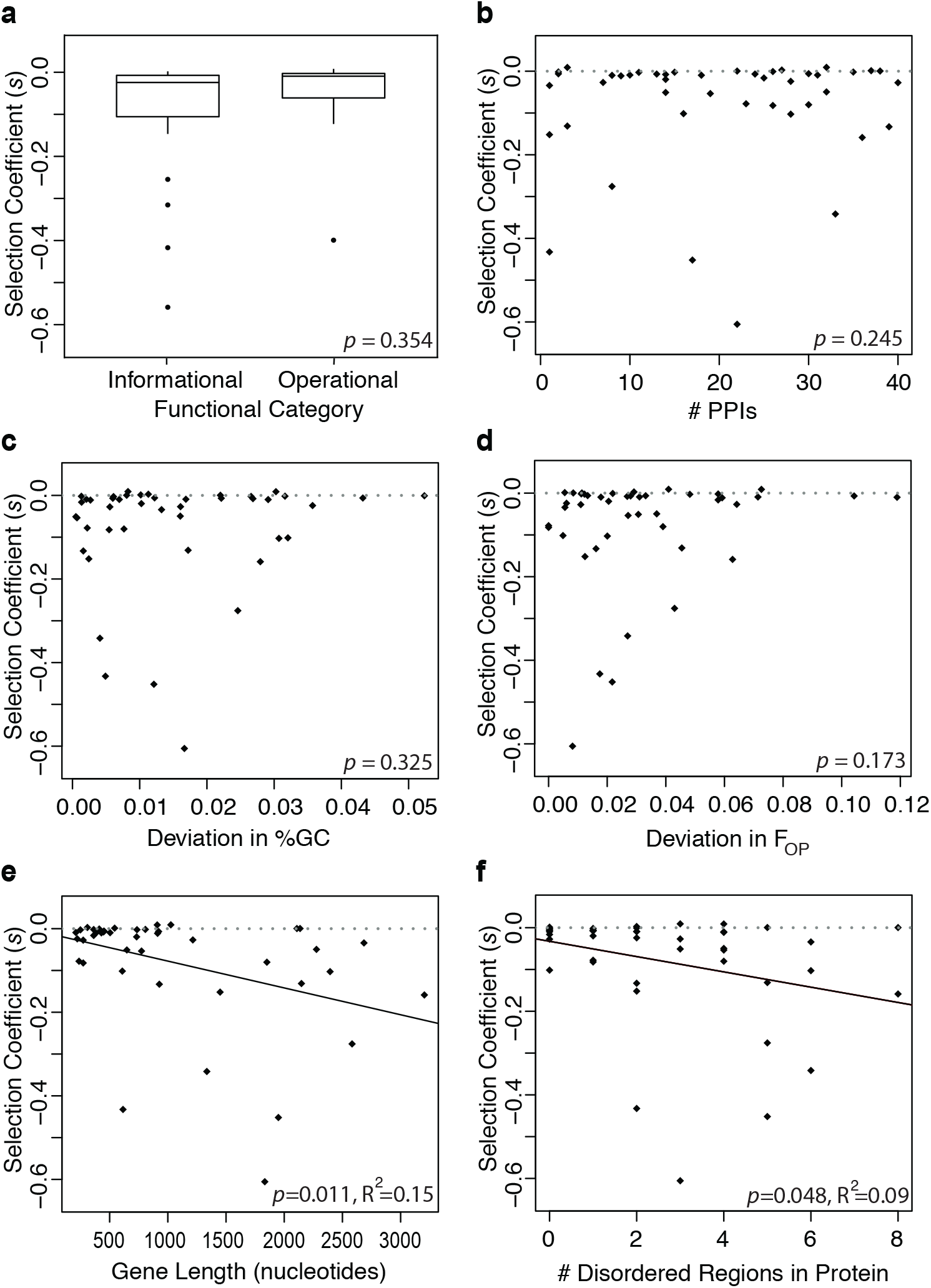
Analyses of the selective barriers on HGT. The mean selective effects of 44 transferred S. Typhimurium genes expressed in E. coli background: **a.** divided into informational and operational genes based on functional categories; **b.** plotted against predicted number of PPIs; **c.** absolute deviation in %GC between orthologs; **d.** absolute deviation in codon usage between orthologs; **e.** gene length; **f.** number of disordered regions in the amino-acid sequences. All 5 factors given in **a.** to **e.** are used as explanatory variables in a multiple regression model (single p-values at the bottom right of each panel). **e.** We repeated the simple linear regression for the gene length as it was the only factor with a significant effect in the multiple regression. Black lines in **e** and **f** show the best fit for the simple linear regression between the two variables; gray dashed lines show the zero line.

### Protein-protein interactions

The ‘complexity hypothesis’ (Cohen et al., 2011) proposes that newly acquired proteins may form spurious interactions with the proteins already present in the recipient cell because of the epistatic incompatibilities accrued during divergence. In other words, an increase in the number of PPIs may decrease the likelihood of successful HGT. We asked whether the number of PPIs affected the fitness effects of transferred genes. We selected 44 genes that represent a range of PPI levels from 1 to 40, covering 83% of the range of *E. coli* genes given in the database of Hu *et al*. (Hu et al., 2009) (see Materials and Methods — Selection of genes). We determined whether potential interacting partners are expressed in the recipient cell under our experimental conditions and adjusted the predicted PPI levels to exclude unexpressed genes (see Materials and Methods — RNA-seq). In contrast to the predictions of the complexity hypothesis, we found that the number of PPI were uncorrelated with observed fitness effects (Fig. 3b, multiple regression model, *p* = 0.245).

### GC content and codon usage

Differences in GC content (Lucchini et al., 2006) and codon usage (Navarre et al., 2006) between a donor and a recipient have been proposed as barriers to horizontal acquisition of genes. Horizontally transferred genes with low GC content may avoid deleterious consequences through H-NS mediated gene silencing in gram-negative bacteria (Lucchini et al., 2006). On the other hand, a fitness cost can arise because of non-optimal codon usage of transferred genes, leading to high rates of translation error, creating toxic side products in the recipient cell (Drummond & Wilke, 2009), or leading to ribosomal sequestration that slows down the overall growth of the recipient cell (Gingold & Pilpel, 2011; Roller, Stoddard, & Schmidt, 2016; Shah, Ding, Niemczyk, Kudla, & Plotkin, 2013; Tuller et al., 2011).

To test the effects of GC content and codon usage, F_OP_ (Ikemura, 1981), we calculated the absolute deviations between our 44 *S*. Typhimurium genes and their *E. coli* orthologs. We examined the effects of the absolute deviation in GC content and codon usage bias separately, while also accounting for the potential correlation between them (simple linear regression, *F_1,42_* = 0.008, *p* = 0.93). In terms of the absolute deviation in GC content, the 44 gene pairs cover a range of 0.1 to 5.2%, which spans the range of 94% of all the ortholog pairs between *S*. Typhimurium and *E. coli*. We found that the absolute deviation in GC content between the transferred *S*. Typhimurium genes and their *E. coli* orthologs did not correlated with the observed fitness effects (Fig. 3c, multiple regression model, *p* = 0.325). With respect to absolute deviation in codon usage bias, the 44 gene pairs cover a range of 0 to 12%, spanning the range of 99% of all the ortholog pairs between the donor and recipient. The factor of absolute deviation in codon usage bias was not a significant predictor of the observed fitness effects either (Fig. 3d, multiple regression model: *p* = 0.173).

### Gene length and intrinsic protein disorder

We identified a significant negative relationship between gene length and the fitness effects of transferred genes (Fig. 3e, multiple regression model: *p* = 0.016, simple linear regression: R^2^ = 0.15, *p* = 0.011). The observed effect is unlikely to arise from the costs of DNA replication and protein synthesis, as these costs are negligible for the small differences in length investigated here, with any effect likely to be below the experimental limit of detection (Baltrus, 2013; Lynch & Marinov, 2015).

One explanation is intrinsic protein disorder — inherently unstable or flexible regions of a protein —which are correlated with gene length (simple linear regression, *p* < 0.001, R^2^ = 0.67, Sup. Fig. 3). While little is understood about disordered regions of proteins, such regions potentially give rise to promiscuous molecular interactions, misfolding, and aggregation (Vavouri, Semple, Garcia-Verdugo, & Lehner, 2009). We identified the number of disordered regions within the protein sequences of the 44 transferred genes using Globplot (globplot.embl.de, Sup. Table 2, Linding, Russell, Neduva, & Gibson, 2003), and found that the *S*. Typhimurium orthologs with more disordered regions in their protein sequences have significantly higher fitness costs (Fig. 3f, simple linear regression, *p* = 0.048, R^2^ = 0.090).

### Dosage sensitivity

Gene dosage effect, or the sensitivity to the change in the concentration of a gene product, is another potential barrier to HGT. Transfer events can result in additional copies of the gene within the recipient cell, potentially yielding a change in the protein concentration and lowering fitness by inducing an imbalance in the stoichiometry of the cell (Bershtein et al., 2015; Papp et al., 2003; Park & Zhang, 2012). Sorek et al. (2007) showed that, at least for some genes, an increase in dosage results in toxicity. Accordingly, we asked whether dosage sensitivity contributes to the observed fitness effects of transferred genes. Dosage sensitivity has classically been studied by modulating the number of gene copies, and hence the level of expression, then measuring the effects on the desired phenotype (Birchler & Veitia, 2012). To test whether dosage sensitivity acts as a barrier to HGT, we measured the fitness effects of transferring additional copies of the native *E. coli* orthologs of the 44 genes into *E. coli* background using the same experimental setup (Sup. Fig. 4). Using this design, any observed changes in fitness must arise from an imbalance in cellular protein levels, *i.e*., dosage sensitivity — rather than the effects of divergent function, number of interacting proteins, or codon usage — as the transferred genes were exact copies of existing genes.

We compared the fitness of each *E. coli* gene (grey bars, Fig. 4) with that of their orthologous *S*. Typhimurium gene that showed significant deleterious effects (n = 36, white bars, Fig. 4). If the *E. coli* copy was equal to or more deleterious than the *S*. Typhimurium copy, we attributed dosage sensitivity as the dominant factor determining the fitness effects of the transferred gene. Alternatively, if the *S*. Typhimurium copy is more deleterious, then factors other than dosage sensitivity drive the fitness effects of the gene. For simplicity, we refer to these groups as dosage sensitive and insensitive, respectively.

**Fig. 4.**
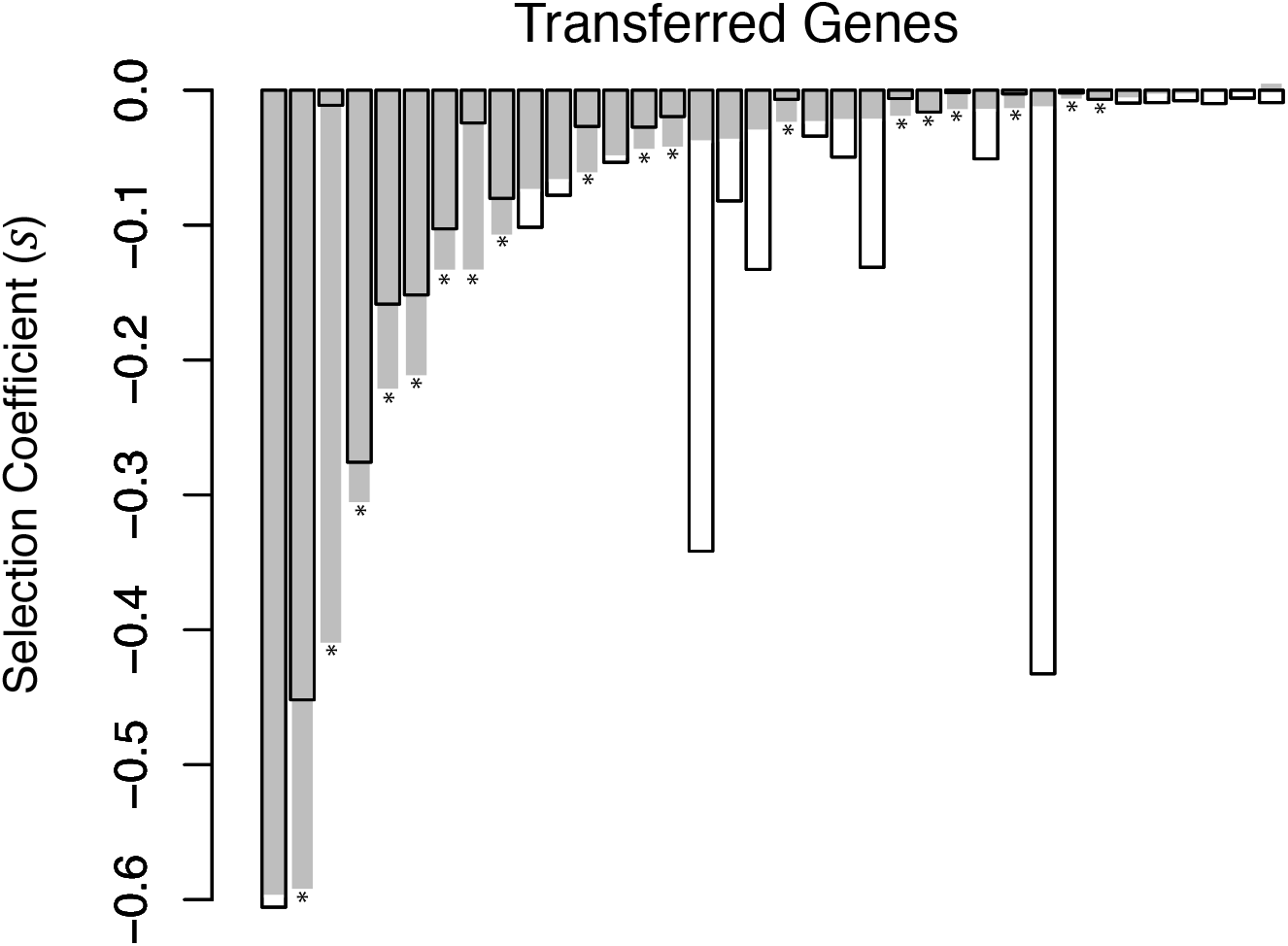
Selection coefficients of newly transferred and deleterious genes from S. Typhimurium (white bars) expressed in E. coli background overlaid with those of their orthologs from E. coli (grey bars) expressed in E. coli background. On the x-axis genes are sorted according to their selection coefficients for the transfer of E. coli orthologs. Genes marked with asterisks show dosage sensitivity, as the fitness cost of the additional copies of an E. coli gene is the same or greater than the fitness cost of its S. Typhimurium ortholog.

For 16 orthologous pairs, the *E. coli* copy was more deleterious than *S*. Typhimurium copy and 2 orthologous pairs had similar fitness effects. Thus 18 out of 36 *S*. Typhimurium genes were dosage sensitive. The remaining 18 genes, where the *S*. Typhimurium copy was significantly more deleterious than the *E. coli* copy, were dosage insensitive (Fig. 4).

We asked whether the previously identified barriers to HGT— functional category, number of PPIs, GC content, codon usage, gene length, and the number of disordered regions — determined the fitness effects of dosage sensitive and dosage insensitive genes independently. For dosage sensitive genes, only gene length (simple linear regression: *p* = 0.002, R^2^ = 0.456) and the number of disordered regions (simple linear regression: *p* = 0.006, R^2^ = 0.391) were significant predictors of fitness (Fig. 5). Surprisingly, these barriers did not predict the fitness effects of dosage insensitive genes (Fig. 5), even though the dosage sensitive and insensitive genes did not differ in their length (Wilcoxon test: *W* = 186, *p* = 0.451, Sup. Fig. 5a) or in the number of disordered regions in their amino-acid sequences (Wilcoxon test: *W* = 145, *p* = 0.602, Sup. Fig. 5b).

**Fig. 5.**
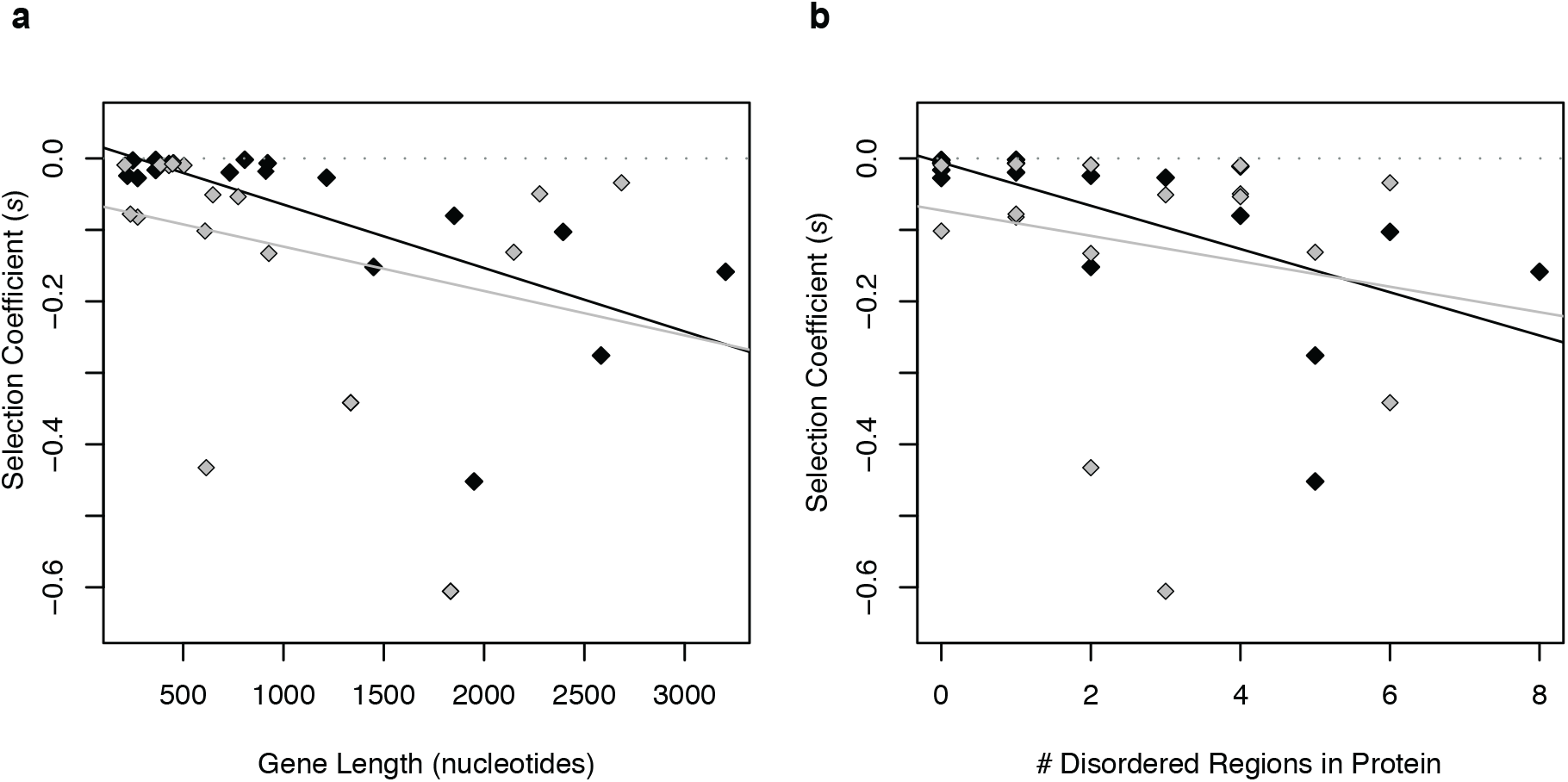
Selection coefficients of 36 newly transferred and deleterious S. Typhimurium genes plotted against **a.** gene length in nucleotides; **b.** number of disordered regions in their amino-acid sequence. Black data points represent the dosage sensitive genes, and grey data points with black frames represent the dosage insensitive genes. Lines are the simple linear regressions between the two variables with corresponding colors, gray dashed lines show the zero line.

Overall, our results suggest that genes with high numbers of disordered regions, and those exhibiting dosage sensitivity, may be less likely to be transferred due to pathological molecular interactions, misfolded toxic configurations, or deleterious aggregates.

## Discussion

Our understanding of HGT has relied primarily on comparative computational analyses of genomes, which can only study successful transfer events that have gone through the sieve of natural selection. Experimental approaches, which have been rare due to their labour-intensive nature and technological limitations, offer the potential to test the role of a different set of factors that could promote or hinder HGT (Knöppel, Lind, Lustig, Näsvall, & Andersson, 2014; Sorek et al., 2007). We relied on a very precise experimental approach for measuring fitness to better understand the role of different selective factors in HGT, which has allowed us to formulate and test new hypotheses about previously unconsidered barriers to HGT.

We explored the relative importance of several previously hypothesized factors that may affect the probability of an HGT event, by transferring orthologous genes from *S*. Typhimurium into *E. coli*. Unlike the mostly neutral effects of transferred DNA fragments of random size, for which the expression state was not known (Knöppel et al., 2014), we find that the majority of transferred genes that are expressed in the recipient cell impose a significant fitness cost. This finding suggests that, if expressed, a large fraction of transferred genes would be rapidly eliminated by selection – a likely scenario given the ease of evolving constitutive promoters in bacteria (Yona et al., 2018). Yet, despite most transfer events resulting in fitness costs, HGT is still one of the major sources of novel genetic material in microbes (Boto, 2010; Soucy et al., 2015). The disparity between this observation and the DFE we report could be reconciled in a couple of ways. Firstly, if the bacterial effective population sizes in nature are smaller than believed, due to spatial structure,recurrent bottlenecks, or recurrent selective sweeps, genetic drift or draft will dominate over selection (Gillespie, 2000; Kimura, 1968). Secondly, the ubiquitous mechanisms for gene silencing may also explain this discrepancy as silencing shields the recipient cell from deleterious effects (Navarre, 2016). Thus, deleterious genes may persist long enough to be rescued by beneficial compensatory mutations or a change in the selective environment.

Interestingly, we do not find evidence for any of the previously hypothesized barriers to HGT: functional gene categories, the number of PPIs, GC content, or optimal codon usage (Baltrus, 2013). With increased divergence between donor and recipient species, these factors may become increasingly important. In our dataset, the relatively modest differences in codon usage and GC content between *S*. Typhimurium and *E. coli* orthologs may prevent us from detecting their role as selective barriers to HGT. Nevertheless, among closely related species these intrinsic properties do not appear to be strong selective barriers.

In contrast, computational studies found limited support for dosage sensitivity as a major barrier to HGT, in part because it is difficult to infer dosage from genomic data (Park & Zhang, 2012). In an experimental study, dosage sensitivity was identified as a barrier along with toxicity for a set of lethal but very divergent genes from a variety of bacterial species (Sorek et al., 2007). Our findings, which in contrast focused on transfer of non-lethal genes between two closely related species, shows that dosage sensitivity is a general barrier to HGT. In addition, the role of dosage sensitivity as a barrier to HGT is consistent with the observation that experimentally altering expression levels of native genes can dramatically reduce fitness (Papp et al., 2003). This effect of dose is related to the intrinsic disorder of proteins, which predicts the fitness effects of dosage sensitive, but not of the dosage insensitive genes. Here we show that gene dosage and regions of protein disorder are significant predictors of the success of HGT and that these are likely to be globally important selective barriers. The short-term evolutionary dynamics of HGT and the barriers that determine the success of a transferred gene differ from their long-term evolutionary dynamics, emphasizing the need for experimental approaches to complement bioinformatic analyses.

## Materials and Methods

### Chromosomal Modifications in the Recipient Strain

To differentiate two cell types, we inserted the fluorescent markers *cfp* and *yfp* under control of the constitutive P_λR_ into the phage p21 attachment site on the *E. coli* MG1655 (DSM18039) chromosome using pAH95 from CRIM system and protocol given previously (Haldimann & Wanner, 2001). Briefly, *E. coli* cells containing pAH121 were electroporated with the pAH95, suspended in SOC without AMP, incubated at 37°C for 1 h and at 42°C for 30 min to get rid of the pAH121, spread onto LB agar plates with kanamycin 10 *μ*g/mL and incubated over night at 37°C. Colonies were streak-purified once non-selectively and then tested for antibiotic resistance for stable integration and loss of the pAH121 and by PCR for single integration of the cassette. The insertion is sequence-verified. The *cfp* gene was derived from an *E. coli* strain used in Elowitz *et al*. (Elowitz, Levine, Siggia, & Swain, 2002), and *yfp* gene is the *Venus* gene derived from the pZS123 used in Cox *et al*. (R. S. Cox, Dunlop, & Elowitz, 2010).

We integrated the repressor protein gene *tetR* that suppresses the expression of transferred genes into the phage λ attachment site on the *E. coli* MG1655 att-p21::(CFP/YFP-Kn^R^) chromosome using the plasmids and protocol given previously (Lutz & Bujard, 1997). pZS4Int plasmid contained *tetR* gene under P_N25_ promoter, spectinomycin resistance gene, and origin of replication pSC101. Briefly, the origin of replication was cut out of the plasmid pZS4Int and the rest of the plasmid was ligated back. *E. coli* cells, containing pLDR8 (Diederich, Rasmussen, & Messer, 1992) were then electroporated with this ligated DNA. Cells were incubated first at 42°C for 2 hours to get rid of pLDR8 and then at 37°C overnight on agar plates supplemented with spectinomycin 50 *μ*g/mL to select resistant clones. Colonies were streak-purified once non-selectively and then tested for antibiotic resistance for stable integration and loss of the pLDR8 and by PCR for single integration of the *tetR* cassette. The insertion is sequence-verified.

### Selection of Genes

*S*. Typhimurium and *E. coli* are genetically similar and share ecological environments (Mugnai et al., 2015; Winfield & Groisman, 2003) As we are interested in the role of protein-protein interactions (PPIs) and functional categories, this similarity ensures that the majority of gene products transferred from *S*. Typhimurium are functional and most of their functional partners exist in the *E. coli* genetic background. Although the exact number of interacting partners may differ in *E. coli*, this difference is expected to be minimal. In addition, the two species are sufficiently divergent to allow us to systematically test the effects of several factors of the introduced genes.

More specifically, *Salmonella enterica* serovar Typhimurium LT2 (DSM18522, Genbank AE006468.1 (McClelland et al., 2001) was used as the gene donor. We excluded genes that are known to be mobile genes, such as phage related proteins, transposable elements or insertion sequences, as well as ribosomal and transfer RNAs that are known to be related to the mobile genes. We applied an arbitrary selection method for the 45 genes as follows. Since a random selection of genes would be biased towards large functional modules, and we expected specific functions of the genes to have an effect on fitness, we ensured that the sampled genes were from different functional modules. In addition, we ensured that the sampling included the widest possible range of PPIs — we sampled uniformly from a range of 1 to 40 physical PPI as reported in a previous study (Hu et al., 2009), and we only selected genes with interactions validated using LCMS and MALDI after SPA tagging of proteins in that study.

### Cloning of Selected Genes

Selected genes were introduced into the recipient *E. coli* cells by transformation of a modified version of the pZS* class of plasmids (Lutz & Bujard, 1997, Sup. Fig. 2). This plasmid is maintained in the recipient cell at 3-4 copies allowing us to keep the expression of the introduced gene at moderate levels. The coding regions of the selected *S*. Typhimurium genes (or their endogenous *E. coli* orthologs) were cloned into the pZS*-HGT plasmids under the control of the inducible promoter P_LtetO-1_ (Lutz & Bujard, 1997). Each gene was cloned at the *Avr*II site at 5’-end to ensure that start codon was located at the exact position relative to the promoter and ribosomal binding site. We failed to clone one gene (STM4381, *ulaR*, transcriptional repressor for the L-ascorbate utilization divergent operon, 756 bps) out of 45 selected genes, therefore, rest of the analyses were performed on the 44 successfully cloned genes.

The plasmids were then transferred into *E. coli* MG1655 att-λ::(tetR-Sp^R^) att-p21::(CFP/YFP-Kn^R^) cells and successful transformants were selected on LB agar with ampicillin 50 *μ*g/mL. After two rounds of streak purification on ‘rich M9 medium’ (1x M9 salts (Sigma-Aldrich, M6030), 1% CAA (Sigma-Aldrich, A2427), 0.4% glucose, 2mM MgSO_4_, 0.1mM CaCl_2_) agar plates with ampicillin 50 *μ*g/mL, single colonies were grown overnight in liquid rich M9 medium with ampicillin 50 *μ*g/mL and stored at −80°C. All the cloned genes are sequence-verified.

Same modified version of the pZS* backbone was used for the experiments of 44 *E. coli* orthologs in the *E. coli* background (DNA synthesized and cloned into our pZS* backbone by Epoch Life Science Inc., Sugar Land, Texas, USA).

### Competition Assays

We performed competition assays using *E. coli* MG1655 att-λ::(tetR-Sp^R^) att-p21::(CFP/YFP-Kn^R^) strains, CFP strain carrying the plasmid pZS*-HGT with the transferred gene (referred to as the ‘mutant’ in this study) while the YFP strain carrying the same plasmid without an insert (referred to as the ‘wild type’ in this study). In total 32 replicate competitions were performed across 4 different days for each gene.

All competition assays were done in ‘rich M9 medium’ with ampicillin 50 *μ*g/mL. We determined the inducer concentration as 5ng/mL anhydrotetracycline (ATC, Sigma-Aldrich, 37919) with a titration assay. For each competition assay, first day, frozen stocks were streaked on rich M9 agar plates. Second day, a colony was picked and grown in rich M9 medium for 16 hours. Third day, overnight cultures were diluted 1000x and grown initially for 60 min, followed by the addition of 5ng/mL ATC to initiate the induction of inserted genes, and then grown for another 60 min, until OD ≃ 0.12. Then the two cell types (wild type and mutant) were mixed at equal ratios and competed with each other for 120 min (~3 generations) in 96-well plates. An initial sample (t_0_) was taken at the beginning of the competition and three more samples (t_1_, t_2_, t_3_) were taken every 40 mins ~ each generation. Mutant to wild type ratio was determined by counting 50,000 cells at each sampling with BD FACSCanto II flow cytometer. Using these four ratios, the fitness costs of genes (*s*) were estimated using the regression model ln(1+*s*) = (ln Rt – lnR_0_)/t, where R is the ratio of mutant to wild type and t is the number of generations (Elena et al., 1998).

By conducting competition assays during the deterministic exponential phase of growth and using time-series data, we were able to detect very small differences in selection coefficients (Δs ≈ 0.002, power analysis, Sup. Fig. 6). We have performed additional tests to account for the fitness effects of different fluorescent markers (see Sup. Mat. Text – Accounting for the fitness effects of fluorescent markers).

### RNA-seq: Sample Preparation

To estimate the expression level of transferred genes in our experiment, we cloned the *mCherry* gene into an empty pZS*-HGT plasmid under the hybrid promoter P_LtetO-1_ as an expression control and used RNA-seq to measure the expression level of *mCherry* gene. Cultures grown overnight in rich M9 medium were diluted 1000x and handled under the same conditions as the competition assays described above. When the OD of the cultures reached to ~0.12, growth was stopped by adding Qiagen RNA protect Bacteria Reagent (cat no. 76506) to 20mL cultures (~6×10^8^ cells). Total RNA was purified with Qiagen Rneasy Mini Kit (cat no. 74104). Quality and integrity of the total RNA samples were checked in Agilent 2100 Bioanalyzer and Agilent RNA 6000 Nano Kit (reorder number 5067-1511). The experiment was performed as 3 biological replicates. Library preparation (RiboZero, NEB), further quality checks and next-generation sequencing (HiSeq2500-v4, SR100 mode) were performed at the VBCF NGS Unit (www.vbcf.ac.at).

### RNA-seq: Data Processing

Sequence reads with an average read quality of >= 34 were retained for further analysis. After quality controls, fastq files were mapped against the *E. coli* MG1655 genome (Genbank U00096.3) using the Bowtie2 aligner using default settings in RSEM (Langmead & Salzberg, 2012). The reference genome was modified *in silico* to contain the chromosomal modifications of *tetR* and fluorescent protein gene cassettes. Expected counts were calculated by using the defaults in RSEM (Li & Dewey, 2011). After between-sample normalization of the counts with the DESeq package of the R statistical software (Anders & Huber, 2010), TPM (transcript per million) values for each gene were calculated and used in further analyses (Li & Dewey, 2011, Sup. Data 1).

RNA-seq data have also served to inspect whether all of the listed interaction partners of the 44 selected genes were expressed under our experimental conditions. After obtaining the expression levels for the whole transcriptome, to determine whether a gene is expressed or not, we needed a threshold level of expression below which a gene would have been eliminated from further analyses. To this end, we inspected the expression levels of genes that are known as being repressed under our experimental conditions, i.e., lactose operon and arabinose regulon genes. Expression level of these genes ranged from 0.5 – 10 TPM in our RNA-seq data. Based on this observation and previous similar studies, we set a threshold value of TPM ≥ 10 (Kröger et al., 2013).

To confirm the reproducibility of our RNA-seq based transcriptomic data, we have performed 3 biological replicates. Correlation coefficients calculated for the expression values of the genes above the TPM_10 threshold between replicates were ≥ 0.99 for all combinations.

### Testing the Protein Expression with Fusion-GFP

To confirm the translational expression of transferred genes in our experiments, we prepared GFP fusion protein cloned at the 3’-end of the genes. We randomly chose a subset of genes, and on each of the corresponding plasmids (pZS*-HGT) we first removed the stop codons of the transferred genes, added a GGSGGS linker and the *GFP* gene without its start codon. In this setting, level of GFP expression indicates the expression level of the transferred genes. Cultures grown overnight in rich M9 medium were diluted 1000x and handled under the same conditions as the competition assays described above. One hour after adding the inducer (5ng/mL ATC), OD_600_ and GFP_540_ emissions were measured using 96 well plate reader, and GFP_540_ readings were normalized with the corresponding OD_600_ readings (Sup. Fig. 1).

### Statistical Analysis

To determine whether the fitness effects of genes were significantly different from zero or not, one-sample t-tests were done for the 32 replicates of each gene, with one-tailed (μ_0_ > 0 for beneficial, μ_0_ < 0 for deleterious genes) or two-tailed (μ_0_ = 0 for neutral genes) settings. α = 0.05 was used as the significance level after Benjamini and Hochberg false discovery rate (BH-FDR) corrections for multiple testing (Benjamini & Hochberg, 1995; Fig. 1). The data for all 32 replicates of 44 orthologous genes of *S*. Typhimurium and *E. coli* are given in Sup. Data 2.

After dividing the 44 orthologous genes from *S*. Typhimurium to *E. coli* into two groups according to their functional categories, one-sided Wilcoxon rank sum test (Mann-Whitney U test) was used to decide if the fitness effects of the informational genes were less than that of the operational genes. Similarly, Levene’s test was used to decide if the variance of the distribution of fitness effects of the informational genes was less than that of the operational genes. Analyses were done on the mean selection coefficients of genes for the 32 replicate measurements (Sup. Data 1, Fig. 3a). Note that, in the experiments in which we transferred the *E. coli* orthologs into the *E. coli* background, there is a significant difference in the means and variances of fitness effects between informational and operational genes (Wilcoxon rank sum test: median ***s_Info_***= −0.039, median ***s_Oper_***= −0.015, *W* = 158, *p* = 0.045, and Levene’s test: 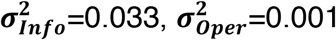, *F_1,41_* = 5.335, *p* = 0.026, respectively).

We investigated a number of intrinsic genetic properties —Protein-protein interaction level (explained under ‘selection of genes’ section), GC content, codon usage, gene length, and disordered amino acid regions. GC content was calculated as the absolute deviation between the introduced *S*. Typhimurium gene and the *E. coli* ortholog. Codon usage was calculated as the absolute deviation of the frequency of optimal (F_OP_) usage in the introduced *S*. Typhimurium sequence using the *E. coli* F_OP_. Gene length was quantified as the number of base pairs from the start to stop codon of the *S*. Typhimurium gene (i.e., cds). We used the web service GlobPlot (http://globplot.embl.de) to identify the number of disordered regions within the protein sequences of the transferred genes from both donors (Linding et al., 2003). To investigate the effect of these intrinsic factors we employed multiple linear regression. After investigating interactions and more complicated models, we used the following model: ‘*S*. Typhimurium selection coefficients’ ~ ‘Functional Category (as dummy variable)’ + ‘Protein-protein interaction level’ + ‘Deviation in GC% between orthologs’ + ‘Deviation in codon usage between orthologs’ + ‘Gene length in nucleotides’. The analysis was done on the mean selection coefficients of genes for the 32 replicate measurements (Fig. 3a, b, c, d, Sup. Table 3). An extended table with all the relevant information is attached in Sup. Table 1 and 2, and Sup. Data 1. As gene length was the only factor with a significant effect in this multiple regression analysis, a simple regression between gene length and *S*. Typhimurium selection coefficients was performed separately (Fig. 3e). Additional simple linear regression was performed between ‘*S*. Typhimurium selection coefficients’ and ‘the number of disordered regions in their amino-acid sequence’ (Fig. 3f).

After dividing the 44 orthologous genes from *S*. Typhimurium to *E. coli* into two groups — 18 genes where dosage is the dominant factor and the remaining 18 genes where it is not, simply by comparing 32 replicates of *S*. Typhimurium selection coefficients to *E. coli* selection coefficients for each ortholog pairs using one-sided t-tests — we performed simple linear regression analyses for the rest of the intrinsic genetic properties (protein-protein interaction level, GC content, codon usage, gene length, and disordered regions in amino-acid sequence) on these two groups separately (Fig. 5). Additional two-sided Wilcoxon rank sum tests (Mann-Whitney U test) were performed to test if these two categories of genes are different in their mean gene length and mean number of disordered regions in amino-acid sequence (Sup. Fig. 5). A final simple linear regression was performed to see the relationship between gene length and the number of disordered regions in amino-acid sequence of *S*. Typhimurium genes (Sup. Fig. 4).

All the statistical analyses were done using the R software package (version 3.1.1).

## Acknowledgments

We thank Katharina M. Pöcher for her guidance during the RNA-seq analysis; John F. Baines, Andrea J. Betancourt and Călin C. Guet for valuable discussions and contributions; Rama P. Bhatia and Jay C. D. Hinton for helpful feedback on the manuscript. This work was supported by the European Research Council FP/2007-2013, ERC grant agreement #648440 to J.P.B.

## Supplementary Materials

### Accounting for the fitness effects of fluorescent markers

As we wished to control for any fitness differences that might be the result of introducing two different fluorescent markers, we compared the fitness of the two ‘wild type’ strains, i.e., strains carrying *cfp* vs *yfp* (both with empty pZS*-HGT plasmids) using the competition assay protocol described in the Materials and Methods section. We detected a small but significant difference between the fitness of CFP strain and YFP strain (*s*_CFP>YFP_ = 0.004, SD = 0.010, *t_(314)_*= 7.118, *p* < 0.001). Therefore, we accounted for this difference in the estimation of selection coefficients of introduced genes during the competition assays. We did that by running a set of competitions between these two ‘wild type’ cells during every competition assay in parallel as a control. Since we did the competition assays in the deterministic phase of the growth under pure haploid selection, this difference in the fitness costs of different fluorescent markers is a constant that we subtracted from the estimated selection coefficient of transferred gene. Such that, each estimation of selection coefficient was corrected for with the fitness difference of the two ‘wild types’ in the control wells of corresponding experiments.

To determine whether the introduced genes might show different fitness effects on the different fluorescent backgrounds (CFP strain and YFP strain) we did a reciprocal introduction by cloning a subset of 8 randomly selected genes out of our 44 *S*. Typhimurium genes into the YFP strain and repeated the competition assays. The regression between the selection coefficients of genes (mean of 32 replicates) in CFP strain and YFP strain was highly significant (*F_1,6_*=117, *p* < 0.001, R^2^ =0.943,slope = 1.007), indicating different fluorescent backgrounds do not interact with this subset of genes, and all measured effects are solely due to the introduced genes.

**Sup. Fig. 1.**
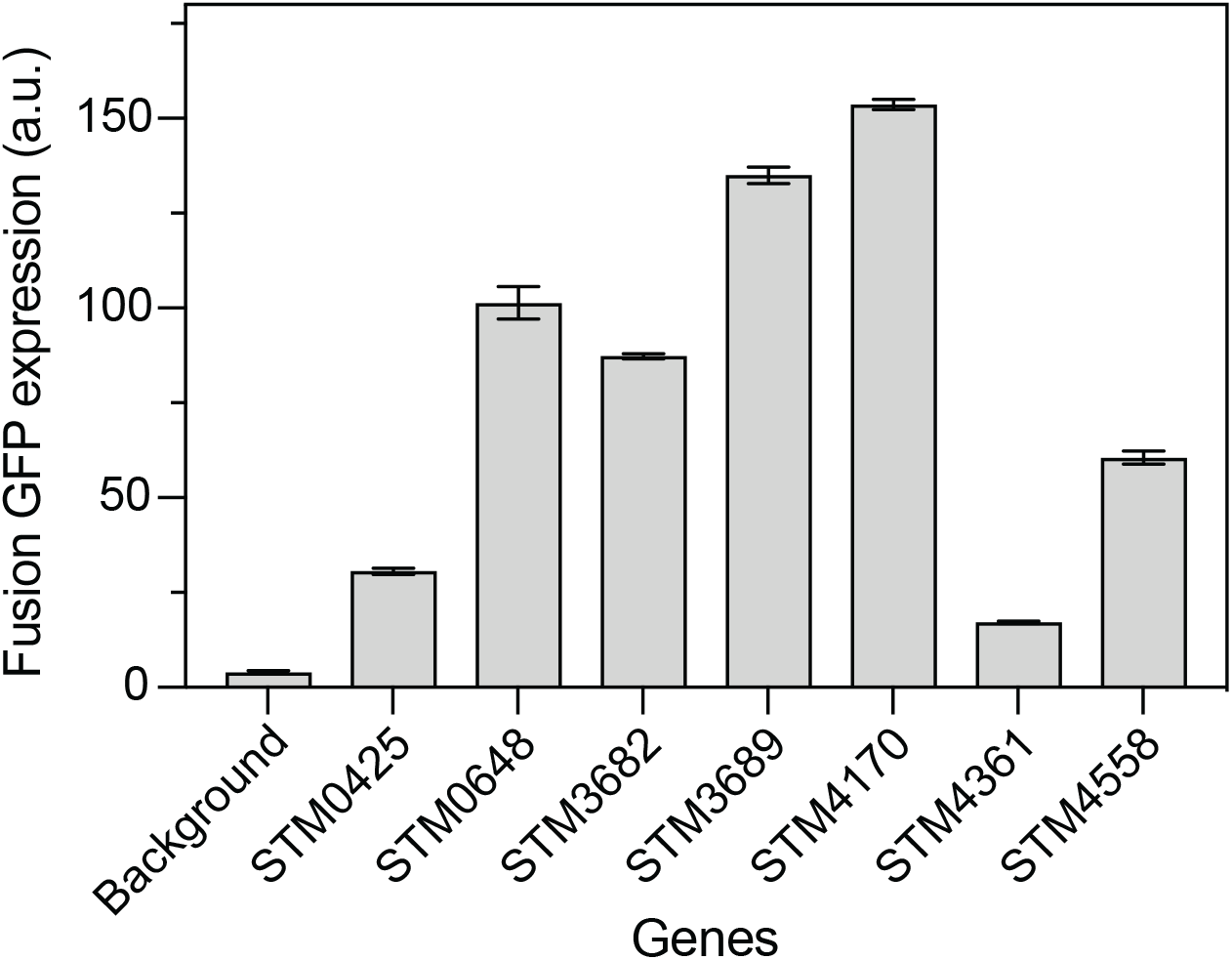
Verifying the expression of a subset of transferred genes at the level of translation. We cloned the GFP gene at the end of the transferred gene as translational fusions. In this setting, level of GFP expression indicates the expression level of the transferred genes. Cultures grown overnight in rich M9 medium were diluted 1000x and handled under the same conditions as the competition assays. One hour after adding the inducer (5ng/mL ATC), OD_600_ and GFP_540_ emissions were measured, and GFP_540_ readings were normalized with the corresponding OD_600_ readings, given here as the Fusion GFP expression. Background is the normalized GFP expression of the E. coli MG1655 att-λ::(tetR-Sp^R^) att-p21::(CFP-Kn^R^) recipient strain carrying the empty plasmid. Error bars represent SD of four replica experiments.

**Sup. Fig. 2.**
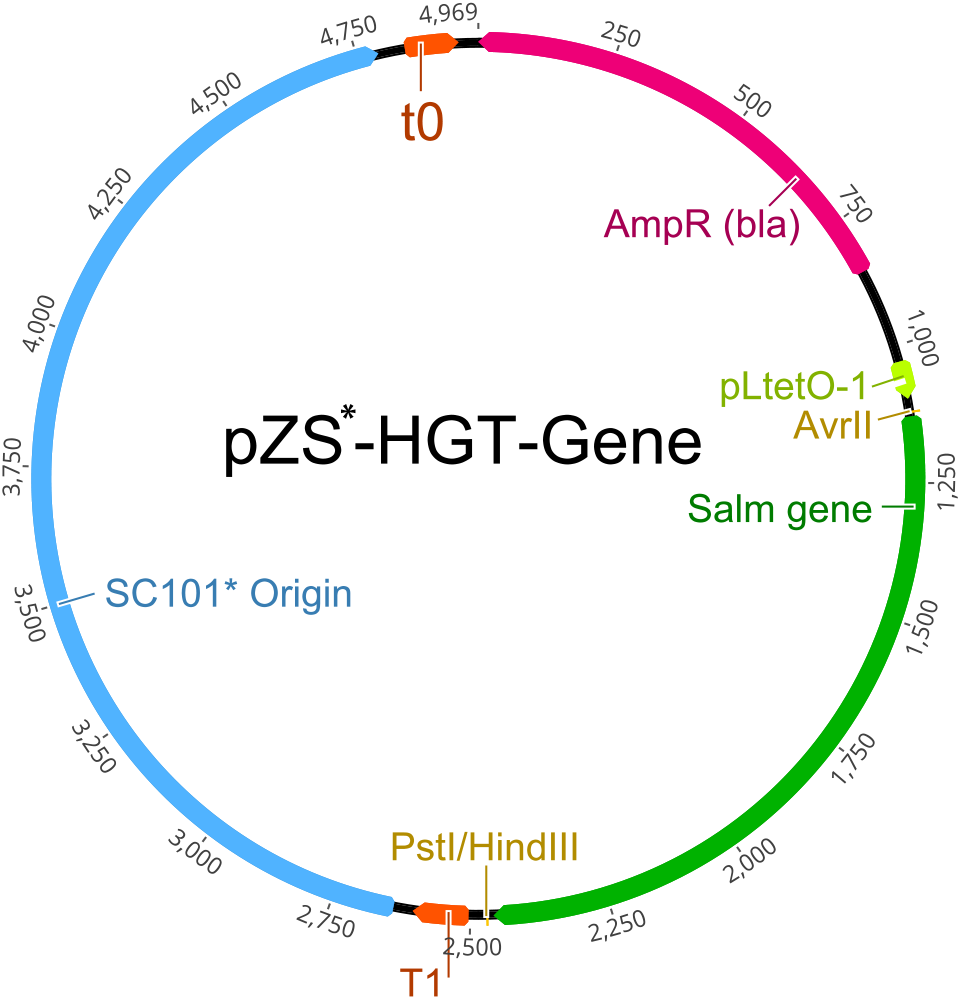
Diagram of the expression plasmid used in this study, modified from Lutz and Bujard (1997). The gene of interest is cloned between the restriction enzyme sites AvrII and PstI/HindIII. The empty plasmid contained only the ribosomal binding site and restriction enzymes sites between the P_LtetO-1_ promoter and the terminator T1. Amp^R^ is the resistance gene for AMP. The terminators T1 of the rrnB operon and t0 of phage lambda.

**Sup. Fig. 3.**
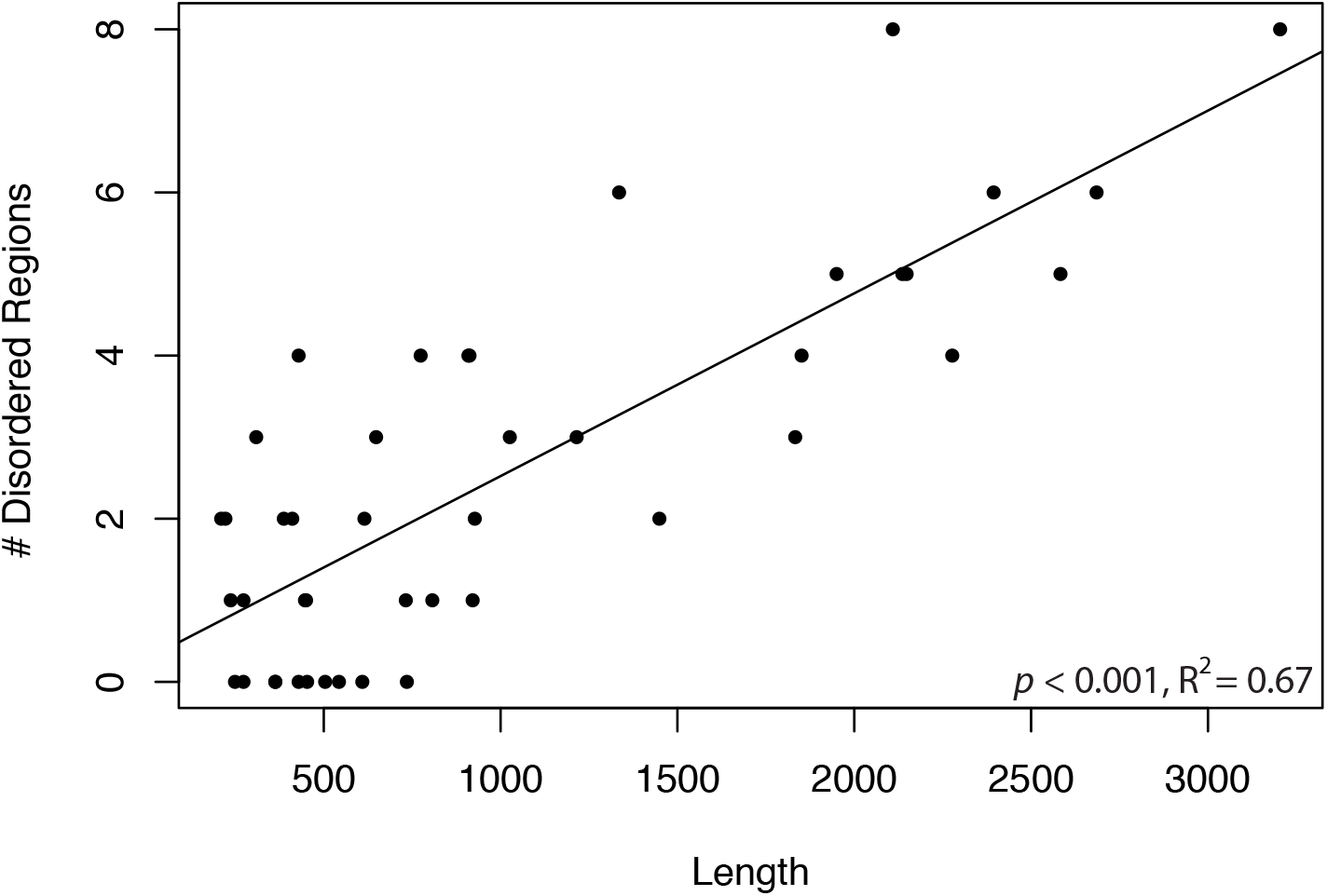
Number of disordered regions in the amino-acid sequences of newly transferred 44 S. Typhimurium genes plotted against gene length (nucleotides). Using simple linear regression, the relationship is significant with p<0.001 (see Materials and Methods – Statistical analyses).

**Sup. Fig. 4.**
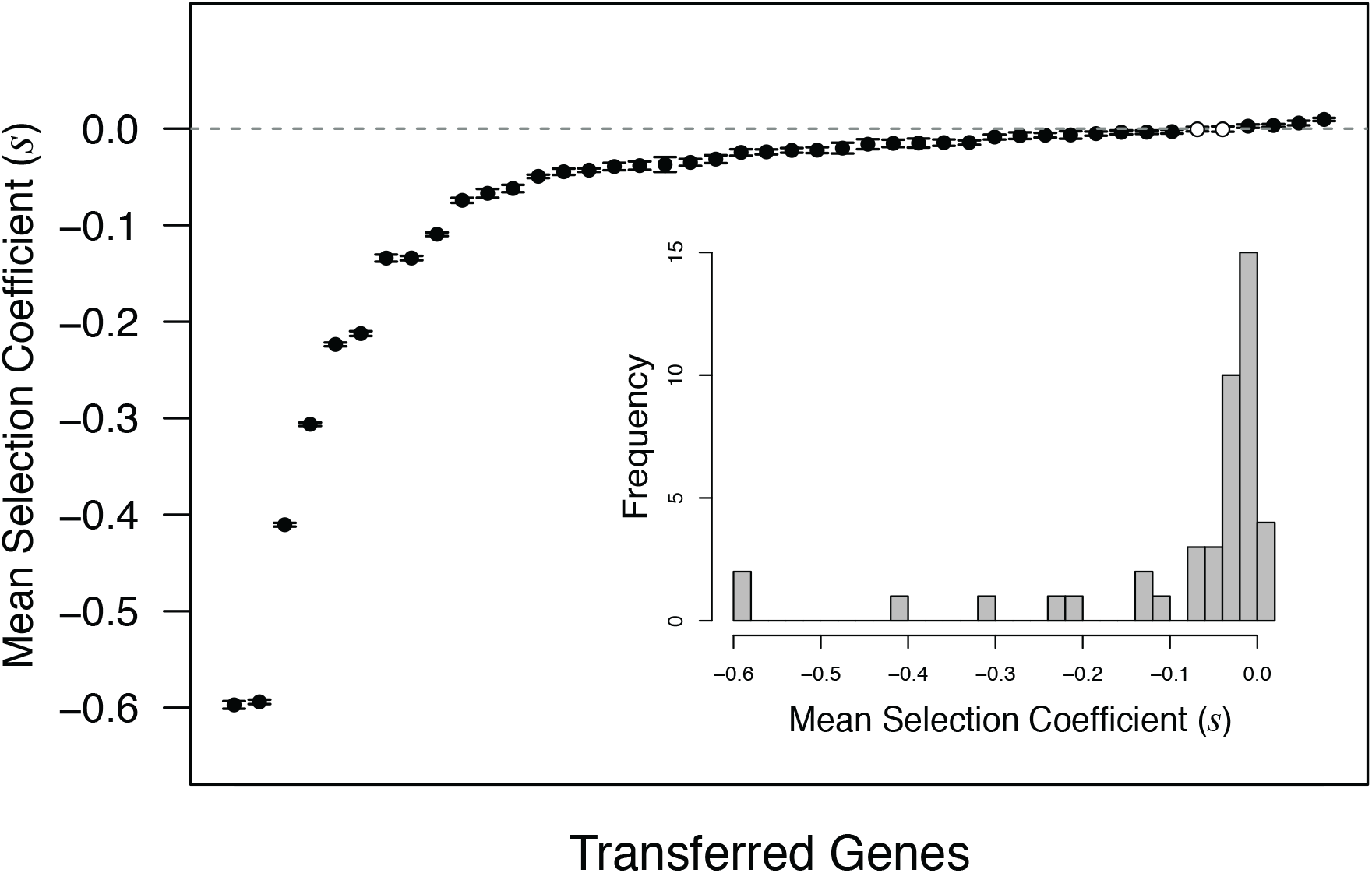
DFE of newly transferred 44 E. coli orthologs expressed in E. coli. On the x-axis, genes are sorted by their fitness effects (s). Error bars indicate the 95% CI of the selection coefficients from 32 replicate measurements for each gene. Empty circles represent the 2 genes with effects not significantly different from zero. Embedded plot gives the histogram representation of the same data.

**Sup. Fig. 5.**
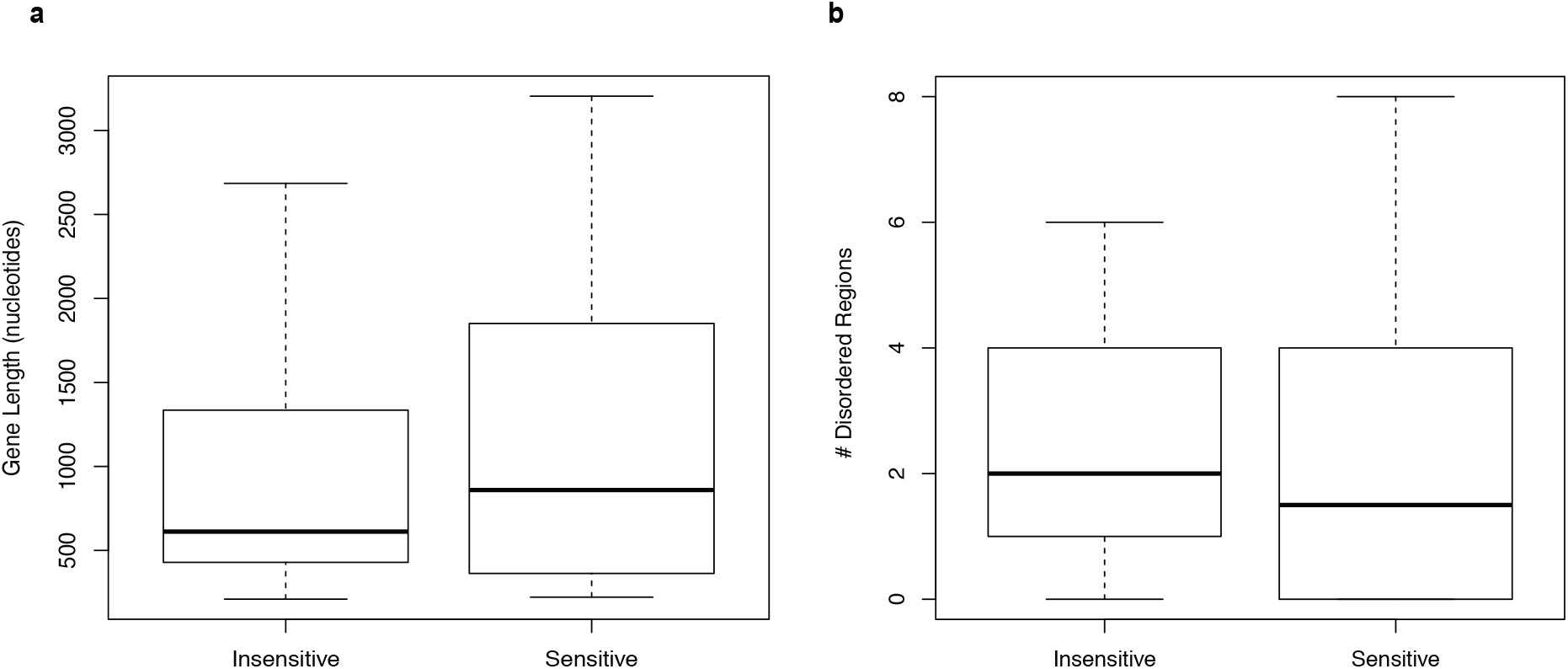
Comparison of 36 newly transferred and deleterious S. Typhimurium genes grouped as dosage sensitive and insensitive. Dosage sensitive and insensitive genes do not differ **a.** in their mean gene length; and **b.** in the mean number of disordered regions in their amino-acid sequences.

**Sup. Fig. 6.**
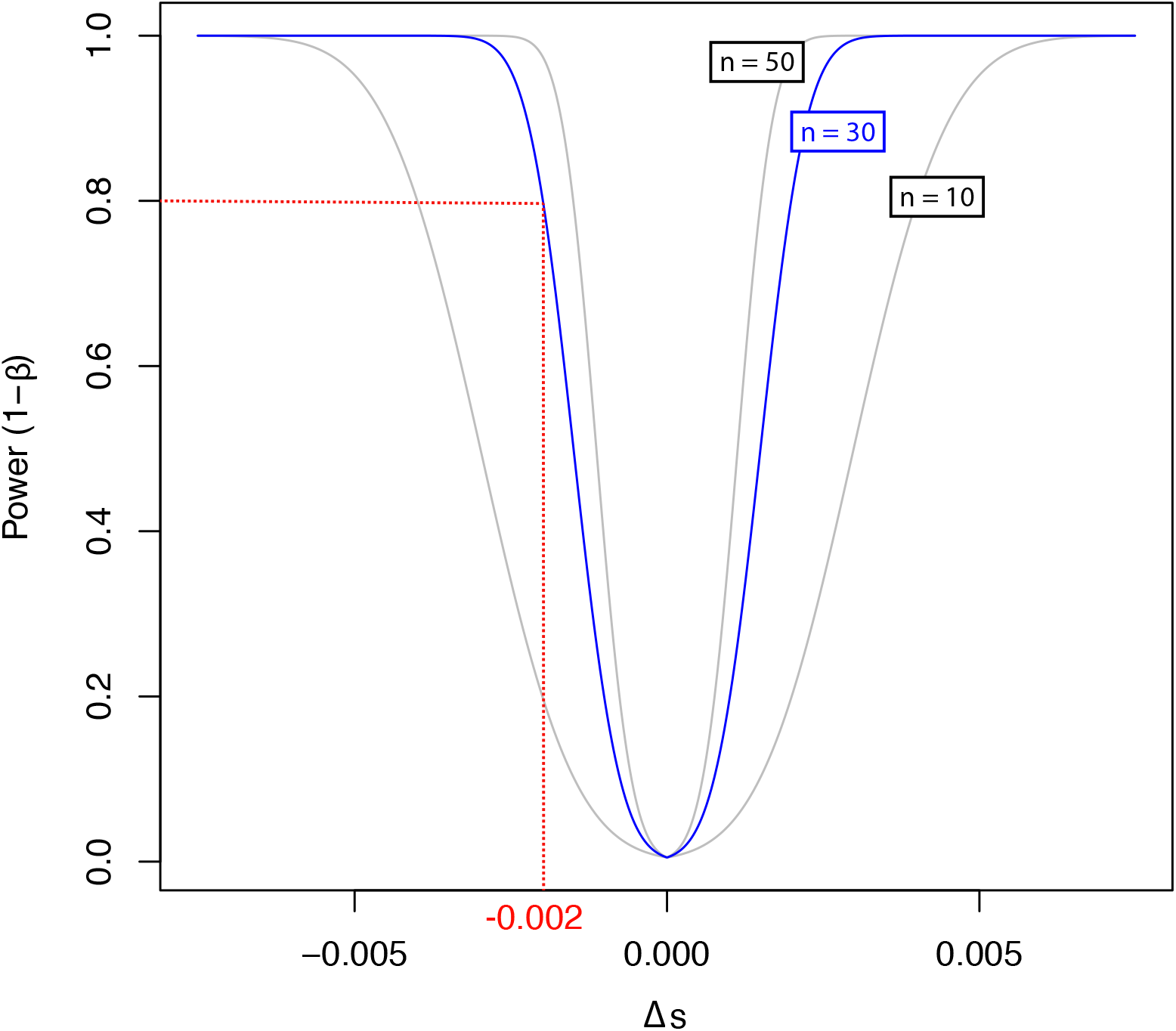
Power analysis performed to estimate the sensitivity of our selection coefficients (s) measurements during competition assays. Power is calculated for α=0.01 and sd=0.003 based on preliminary data.

**Sup. Table 1.** Selection coefficients of all 44 S. Typhimurium and E. coli orthologs expressed in E. coli, sorted according to the selection coefficients of S. Typhimurium orthologs.

| Gene Name | STM Gene ID | STM Sel.Coeff. | p-value | ECO Gene ID | ECO Sel.Coeff. |
| --- | --- | --- | --- | --- | --- |
| *lpxD* | STM0226 | 0.00921 | **<.001** | b0179 | 0.00959 |
| *ybhK* | STM0801 | 0.00896 | **<.001** | b0780 | -0.03490 |
| *hfq* | STM4361 | 0.00285 | **.002** | b4172 | 0.00336 |
| *infC* | STM1334 | 0.00125 | .074 | b1718 | 0.00266 |
| *pnp* | STM3282 | 0.00037 | .719 | b3164 | -0.03917 |
| *rlmL* | STM1061 | 0.00035 | .326 | b0948 | -0.00506 |
| *ydil* | STM1366 | -0.00082 | .102 | b1686 | -0.00670 |
| *sapF* | STM1696 | -0.00181 | **.003** | b1290 | -0.01514 |
| *yacL* | STM0160 | -0.00206 | **.017** | b0119 | -0.00860 |
| *exbB* | STM3159 | -0.00244 | .077 | b3006 | -0.00302 |
| *hybG* | STM3143 | -0.00283 | **.004** | b2990 | -0.01440 |
| *rplI* | STM4394 | -0.00591 | **<.001** | b4203 | -0.00038 |
| *moaE* | STM0806 | -0.00624 | **<.001** | b0785 | -0.02008 |
| *cbpA* | STM1112 | -0.00678 | **<.001** | b1000 | -0.02457 |
| *uspG* | STM0614 | -0.00681 | **<.001** | b0607 | -0.00716 |
| *rimI* | STM4558 | -0.00776 | **<.001** | b4373 | -0.00356 |
| *ridA* | STM4458 | -0.00917 | **<.001** | b4243 | -0.00369 |
| *cspE* | STM0629 | -0.00923 | **<.001** | b0623 | 0.00589 |
| *dps* | STM0831 | -0.00963 | **<.001** | b0812 | -0.00648 |
| *ibpB* | STM3808 | -0.00992 | **<.001** | b3686 | -0.00045 |
| *glyQ* | STM3656 | -0.01117 | **<.001** | b3560 | -0.41062 |
| *yibL* | STM3689 | -0.01623 | **<.001** | b3602 | -0.01602 |
| *dnaQ* | STM0264 | -0.01959 | **<.001** | b0215 | -0.04294 |
| *cspD* | STM0943 | -0.02427 | **<.001** | b0880 | -0.13406 |
| *iscS* | STM2543 | -0.02693 | **<.001** | b2530 | -0.06198 |
| *hupA* | STM4170 | -0.02744 | **<.001** | b4000 | -0.04459 |
| *kdpD* | STM0703 | -0.03411 | **<.001** | b0695 | -0.02397 |
| *clpA* | STM0945 | -0.04958 | **<.001** | b0882 | -0.02243 |
| *yqjI* | STM3215 | -0.05091 | **<.001** | b3071 | -0.01484 |
| *pstB* | STM3854 | -0.05355 | **<.001** | b3725 | -0.04937 |
| *acpP* | STM1196 | -0.07789 | **<.001** | b1094 | -0.06697 |
| *selB* | STM3682 | -0.08022 | **<.001** | b3590 | -0.10939 |
| *hupB* | STM0451 | -0.08211 | **<.001** | b0440 | -0.03703 |
| *lexA* | STM4237 | -0.10157 | **<.001** | b4043 | -0.07425 |
| *malP* | STM3514 | -0.10279 | **<.001** | b3417 | -0.13412 |
| *fadJ* | STM2388 | -0.13126 | **<.001** | b2341 | -0.02219 |
| *yadG* | STM0172 | -0.13294 | **<.001** | b0127 | -0.03153 |
| *thiI* | STM0425 | -0.15182 | **<.001** | b0423 | -0.21239 |
| *rne* | STM1185 | -0.15861 | **<.001** | b1084 | -0.22364 |
| *leuS* | STM0648 | -0.27576 | **<.001** | b0642 | -0.30644 |
| *srmB* | STM2643 | -0.34174 | **<.001** | b2576 | -0.03822 |
| *lolA* | STM0961 | -0.43262 | **<.001** | b0891 | -0.01427 |
| *topB* | STM1298 | -0.45193 | **<.001** | b1763 | -0.59428 |
| *uvrC* | STM1946 | -0.60555 | **<.001** | b1913 | -0.59745 |

*p*-values are for the fitness effects of *S. Typhimurium* orthologs being different from zero. Fields indicated with **bold** are significant values, $\alpha = 0.05$ after corrections for multiple testing with BH-FDR method (Benjamini & Hochberg, 1995).

**Sup. Table 2.** Data used in the multiple linear regression analysis, related to the transferred S. Typhimurium genes.

| STM Gene ID | Functional Category | # PPI | Deviation %GC | Deviation F<sub>OP</sub> | Gene Length | Disordered regions | Dosage |
| --- | --- | --- | --- | --- | --- | --- | --- |
| STM0160 | Unknown | 33 | 0.001 | 0.058 | 363 | 3 | Sensitive |
| STM0172 | Operational | 39 | 0.002 | 0.016 | 927 | 5 | Insensitive |
| STM0226 | Operational | 30 | 0.008 | 0.041 | 1026 | 2 | - |
| STM0264 | Informational | 12 | 0.010 | 0.020 | 732 | 6 | Sensitive |
| STM0425 | Informational | 1 | 0.002 | 0.012 | 1449 | 5 | Sensitive |
| STM0451 | Informational | 20 | 0.005 | 0.000 | 273 | 8 | Insensitive |
| STM0614 | Operational | 12 | 0.022 | 0.028 | 429 | 2 | Sensitive |
| STM0629 | Informational | 31 | 0.002 | 0.071 | 210 | 2 | Insensitive |
| STM0648 | Informational | 8 | 0.025 | 0.043 | 2583 | 5 | Sensitive |
| STM0703 | Operational | 1 | 0.013 | 0.006 | 2685 | 6 | Insensitive |
| STM0801 | Operational | 3 | 0.030 | 0.073 | 909 | 0 | - |
| STM0806 | Operational | 0 | 0.043 | 0.033 | 453 | 1 | Sensitive |
| STM0831 | Operational | 12 | 0.007 | 0.018 | 504 | 4 | Insensitive |
| STM0943 | Informational | 28 | 0.036 | 0.006 | 222 | 1 | Sensitive |
| STM0945 | Informational | 31 | 0.016 | 0.037 | 2277 | 4 | Insensitive |
| STM0961 | Operational | 1 | 0.005 | 0.017 | 615 | 3 | Insensitive |
| STM1061 | Informational | 21 | 0.008 | 0.011 | 2109 | 4 | - |
| STM1112 | Informational | 23 | 0.006 | 0.104 | 921 | 6 | Sensitive |
| STM1185 | Informational | 35 | 0.028 | 0.063 | 3204 | 0 | Sensitive |
| STM1196 | Operational | 20 | 0.002 | 0.000 | 237 | 3 | Insensitive |
| STM1298 | Informational | 16 | 0.012 | 0.022 | 1950 | 2 | Sensitive |
| STM1334 | Informational | 36 | 0.010 | 0.006 | 543 | 1 | - |
| STM1366 | Operational | 25 | 0.052 | 0.022 | 411 | 0 | - |
| STM1696 | Operational | 2 | 0.032 | 0.011 | 807 | 4 | Sensitive |
| STM1946 | Informational | 21 | 0.017 | 0.008 | 1833 | 4 | Insensitive |
| STM2388 | Operational | 3 | 0.017 | 0.045 | 2148 | 0 | Insensitive |
| STM2543 | Operational | 5 | 0.016 | 0.064 | 1215 | 2 | Sensitive |
| STM2643 | Informational | 31 | 0.004 | 0.027 | 1335 | 2 | Insensitive |
| STM3143 | Operational | 7 | 0.006 | 0.048 | 249 | 1 | Sensitive |
| STM3159 | Operational | 14 | 0.027 | 0.012 | 735 | 0 | - |
| STM3215 | Informational | 14 | 0.001 | 0.031 | 648 | 1 | Insensitive |
| STM3282 | Informational | 37 | 0.022 | 0.008 | 2136 | 0 | - |
| STM3514 | Operational | 25 | 0.031 | 0.020 | 2394 | 1 | Sensitive |
| STM3656 | Informational | 9 | 0.003 | 0.059 | 912 | 0 | Sensitive |
| STM3682 | Informational | 29 | 0.008 | 0.039 | 1851 | 0 | Sensitive |
| STM3689 | Informational | 24 | 0.001 | 0.058 | 363 | 0 | Sensitive |
| STM3808 | Operational | 3 | 0.029 | 0.119 | 429 | 1 | Insensitive |
| STM3854 | Operational | 17 | 0.001 | 0.027 | 774 | 2 | Insensitive |
| STM4170 | Informational | 34 | 0.006 | 0.011 | 273 | 8 | Sensitive |
| STM4237 | Informational | 14 | 0.032 | 0.005 | 609 | 5 | Insensitive |
| STM4361 | Informational | 24 | 0.011 | 0.029 | 309 | 0 | - |
| STM4394 | Informational | 27 | 0.012 | 0.013 | 450 | 2 | Insensitive |
| STM4458 | Operational | 9 | 0.017 | 0.031 | 387 | 4 | Insensitive |
| STM4558 | Informational | 12 | 0.027 | 0.027 | 447 | 3 | Insensitive |

**Sup. Table 3.** Result of the multiple regression analysis, F_5,37_ = 2.24.

|  | **R<sup>2</sup>** | **B** | **SE'B** | **beta** | **p-value** |
| --- | --- | --- | --- | --- | --- |
| Model | 0.232 | - | - | - | 0.071 |
| Intercept |  | -0.136 | 0.070 | - | 0.059 |
| Functional Category |  | 0.043 | 0.045 | 0.155 | 0.354 |
| Protein Interactions |  | 2.214e<sup>-03</sup> | 1.874e<sup>-03</sup> | 0.191 | 0.245 |
| Deviation in F<sub>OP</sub> |  | 1.054 | 0.758 | 0.203 | 0.173 |
| Deviation in %GC |  | 1.635 | 1.638 | 0.150 | 0.325 |
| Gene Length |  | -6.278e<sup>-05</sup> | 2.489e<sup>-05</sup> | -0.370 | 0.016 |

